# CyclicBoltz1, fast and accurately predicting structures of cyclic peptides and complexes containing non-canonical amino acids using AlphaFold 3 Framework

**DOI:** 10.1101/2025.02.11.637752

**Authors:** Xuezhi Xie, Christina Z Li, Jin Sub Lee, Philip M Kim

## Abstract

Cyclic peptides exhibit favourable properties making them promising candidates as therapeutics. While the design and modeling of peptides in general has seen rapid advances since the advent of modern machine learning methods, existing deep learning models cannot effectively predict cyclic peptide structures containing non-canonical amino acids (ncAAs), which are often crucial for peptide therapeutics. To address this limitation, we here extend the recent AF3-style model Boltz to cyclic peptides with ncAAs. In addition to the positional encoding offset, we used a simple yet effective extension of the cyclic offset encoding based on AlphaFold3’s tokenization scheme that allows the modeling of cyclic peptides with modified residues. On a test set of peptides with ncAAs, our approach outperforms HighFold2 on 13/17 cases, with an average C*α* RMSD of 1.877Å and an average all-atom RMSD of 3.361Å. Our results show that the cyclic offset encoding shown for AlphaFold2 generalizes to AlphaFold3-based models and can be extended to incorporate ncAAs, showing potential in the design of novel cyclic peptides with ncAAs for therapeutic applications.

## Introduction

Cyclic peptides have emerged as a crucial class of therapeutics due to their enhanced stability, membrane permeability, and specificity compared to linear peptides. Unlike linear peptides, which are rapidly degraded by proteases and exhibit poor bioavailability, cyclic peptides adopt constrained conformations that improve their resistance to enzymatic degradation and enhance their ability to target protein-protein interactions with high affinity, enabling them to be great therapeutic candidates [3]. According to a report published in *Pharmaceuticals* (2023) [15], a total of 53 cyclic peptides have received regulatory approval, accounting for 46% of all approved peptide therapeutics. This highlights their clinical significance and the rapid growth of cyclic peptide-based therapeutics. Cyclic peptides often incorporate engineered non-canonical amino acids (ncAAs), further improving their pharmacological properties by increasing structural rigidity and reducing proteolytic cleavage [17]. However, designing and modeling cyclic peptides remain challenging due to limited structural data, where the Protein Data Bank (PDB) [13] only contains a few hundred cyclic protein or peptide structures, hindering effective modeling of these peptides. Computational tools have shown promise in predicting the structures of cyclic peptides composed of natural amino acids [18][19][20], but their ability to model cyclic peptides with ncAAs remains limited, in particular for machine learning methods.

Traditional cyclic peptide design approaches, such as heuristic computational modeling methods or high-throughput experimental screening, have shown success, but are often very resource-intensive and inefficient [18][19][20]. State-of-the-art deep learning models, such as AlphaFold3 have enabled both protein structure and complex predictions [4][5][6]. Due to the dearth of structural data for cyclic peptides, there has been a focus on modifying pre-trained protein structure prediction models such as AlphaFold2 [7] and RoseTTAFold [8] to cyclize the generated peptide structures. The AfCycDesign model has previously shown that cyclic constraints can be enforced by applying a cyclic offset encoding to existing relative positional encodings in AlphaFold2 to generate cyclic peptide monomers [9]. A similar cyclic offset encoding has been employed to predict cyclic peptide-protein complexes, as demonstrated by HighFold [10], which modifies AlphaFold2-Multimer, and RFpeptides [11], which utilizes RFdiffusion [12].

Despite these advances, current models are largely restricted to generating cyclic peptides composed exclusively of standard amino acids. Concurrently with our work, HighFold2 [16] —an extension of AlphaFold2-Multimer—has been fine-tuned on linear peptides incorporating a limited set of unnatural amino acids. However, this approach remains constrained by the size of the training dataset and the underlying framework, supporting only a few unnatural amino acids and requiring energy functions for post-optimization. The recent release of Boltz1, an open-source implementation of AlphaFold3, presents a new opportunity to address these limitations, since AlphaFold3 can model diverse biomolecular entities including not only proteins and modified amino acids, but also nucleic acids, small molecule ligands, and ions. We hypothesized a similar cyclic offset encoding approach would be effective for the prediction of cyclic peptides with ncAAs.

Building on advancements in the AlphaFold3 framework, we introduce CyclicBoltz1, a method for encoding cyclization and assessing the structural accuracy of cyclic peptide monomers and complexes derived from the Protein Data Bank (PDB) [13]. Previous cyclic offset encoding strategies proved ineffective for Boltz1 since the tokenization scheme of AlphaFold3 treats standard residues as a single token, but ncAAs as atom-wise tokens. To address this limitation, we developed a modified cyclic offset encoding matrix that ensures all atoms in a single ncAA residue are considered a part of one residue (Supplementary Algorithm 1). Our approach outperforms both AfCycDesign [9] and HighFold [10], demonstrating superior accuracy in predicting the structures of cyclic peptides, including those containing noncanonical amino acids (ncAAs). Notably, CyclicBoltz1 achieves a lower average all-atom RMSD of 3.361 Å, compared to 4.097 Å with HighFold2 [16], thereby enabling the generation of a diverse library of structured macrocycles. Furthermore, to facilitate peptide binder design, we propose a cyclic peptide design pipeline tailored for specific targets.

## Results

### CyclicBoltz1, predicting the structures of cyclic peptides monomers and complexes

The CyclicBoltz1 framework introduces two specialized modules — CyclicBoltz1-monomer and CyclicBoltz1-multimer — to predict cyclic peptide structures, including those with ncAAs. At its core, the method modifies the relative positional encoding matrix within Boltz1 to enforce cyclic constraints. During structure prediction, the model generates multiple sequence alignments (MSAs) and pairwise features using Boltz1’s existing MSA generation process, but replaces the standard residue index offset matrix with a custom cyclic offset matrix that accounts for the atom token representation used for modified residues (Supplementary Algorithm 1). This cyclic offset matrix adjusts the sequence separation between terminal residues for peptides, setting it to one, and encodes circularization into the relative positional matrix. These features are passed to the Boltz1 module, where the cyclic positional encoding is incorporated into features to maintain order invariance and enforce cyclic topology. These modifications were implemented within the Boltz-1 framework, enabling robust cyclic structure prediction and design. Initial tests using sequences from the Protein Data Bank (PDB) confirmed that the cyclic constraints produced correct peptide bond connections and preserved terminal residue geometry without distorting the remaining peptide structure. Furthermore, predictions for circularly permuted sequences consistently generated near-identical structures, showcasing the model’s invariance to sequence order. Additionally, we propose a novel pipeline (Fig. 1C) integrating protein structure prediction and *in silico* directed evolution to design binders. This framework optimizes peptide binder sequences by minimizing hotspot distances and maximizing predicted confidence.

**Figure 1.**
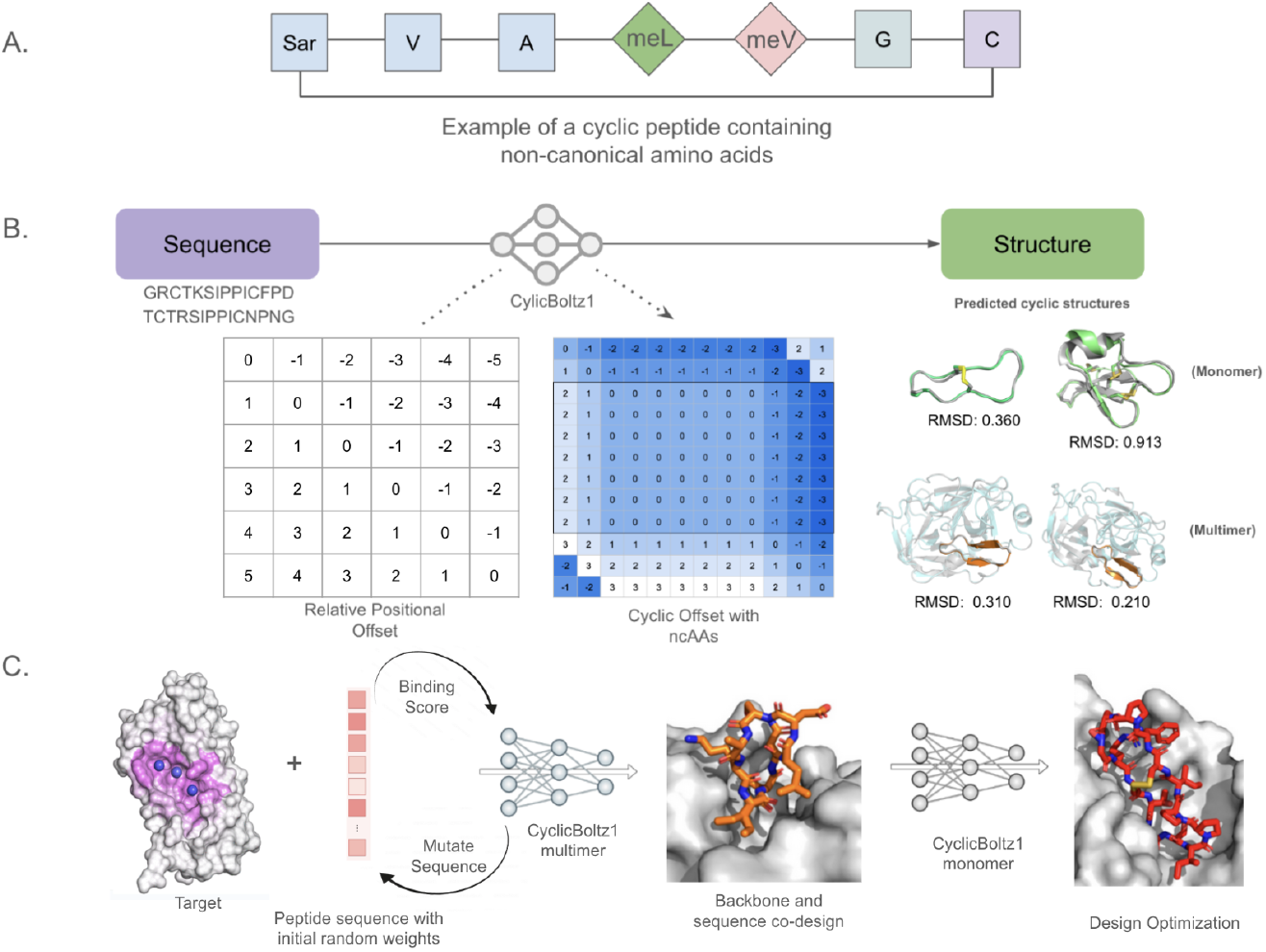
The workflow of CyclicBoltz1. **A**. One example of a cyclic peptide containing ncAAs. The sequence shown uses HELM representation. **B**. CyclicBoltz1 encompasses a duet of modules: CyclicBoltz1-monomer and CyclicBoltz1-multimer, and works for ncAAs. The core idea is to update the relative positional encoding matrix inside Boltz1 to cyclize the generated structures. When given an input sequence, the model first produces multiple sequence alignments (MSAs) and pairwise features. The construction of MSAs follows that of the original Boltz1, but the original residue_index offset matrix is replaced with a cyclic offset matrix, enabling the module to predict corresponding structures. **C**. Proposed schematic representation of the CyclicBoltz1 binder design pipeline. The hotspot residues are highlighted in purple. Given a target protein structure, a binder backbone and sequence are generated using CyclicBoltz1-multimer, which are then filtered using the CyclicBoltz1-monomer model.

**Figure 2.**
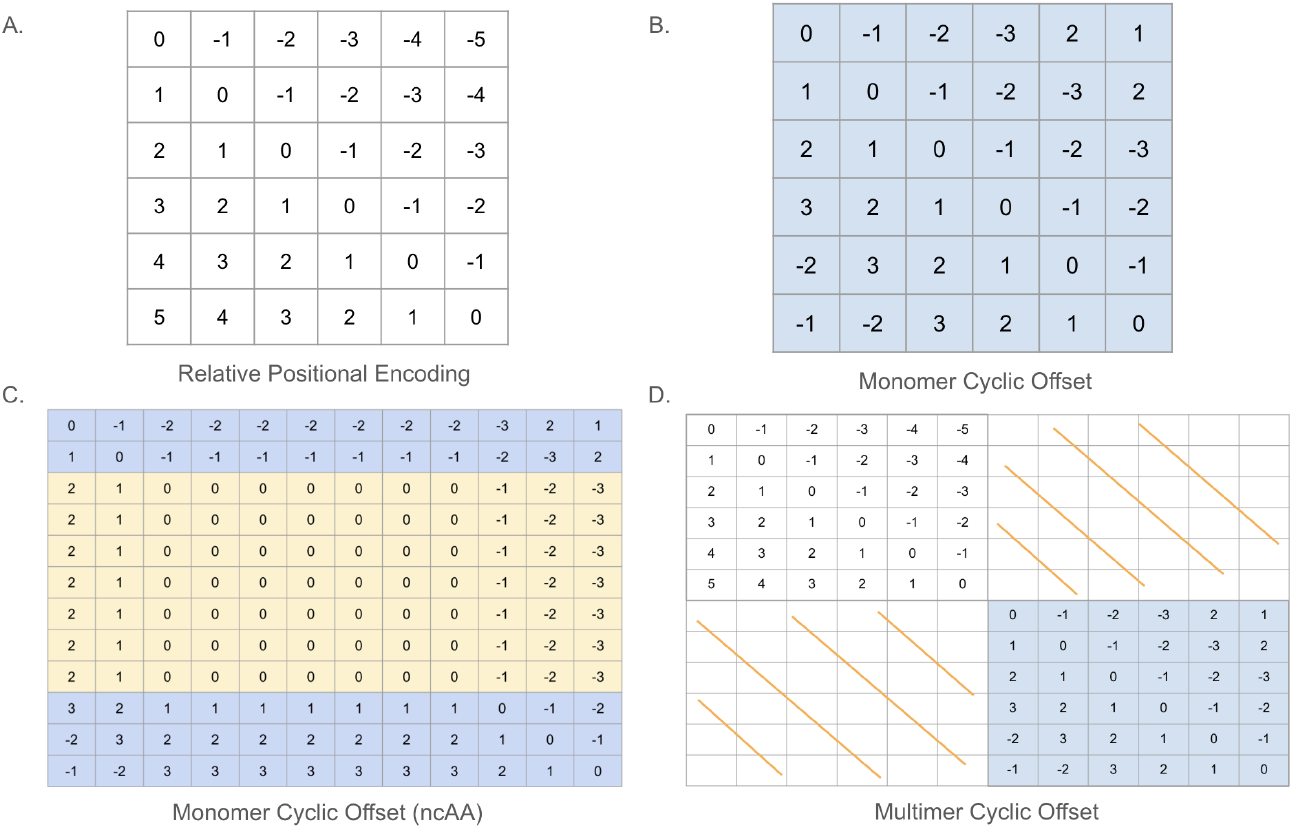
Relative encoding illustrations. **A**. Original relative positional encoding in Boltz1. **B**. The updated matrix in peptide monomers with standard amino acids. **C**. The updated matrix in CyclicBoltz1-monomer with a ncAA inserted at residue 3, where the section highlighted in yellow are seven atoms of a ncAA. **D**. The cyclic positional encoding matrix for multimers.

### Prediction of Cyclic Peptide Monomers

We tested the CyclicBoltz1 model that contains different cyclic offset matrices for predicting the structures of diverse cyclic peptides under different experimental settings (monomer vs multimer predictions). From the publicly available Highfold dataset, we tested all 63 monomeric cyclic peptide structures. The test set was chosen due to that both HighFold (AlphaFold 2.3) and Boltz share the same training data cutoff dates. Using the CyclicBoltz1 monomer, we predict the three-dimensional structures for each sequence in the dataset. We then assessed the backbone C*α* RMSD as a metric to evaluate against the corresponding experimentally determined ground truth structures.

We found that the CyclicBoltz1-monomer model had better performance compared to existing models. As our control, we used the reported C*α* RMSD for the corresponding test cases for the AfCycDesign and HighFold models from the HighFold paper. Since they reported the values for the top-1 prediction, we used the top-1 prediction from the samples generated by our model for comparison. Among three methods, we found that our model showed lower C*α* RMSD in 32 out of 63 test cases, demonstrating superior performance on a significant subset of the test set (Fig. 3A). As expected, we noticed a negative correlation between the C*α* RMSD and the pLDDT indicating that the higher RMSD values were likely due to poor confidence predictions for the cyclic peptide monomers and also noticed a positive correlation between the C*α* RMSD and the peptide length (Fig. 3B-C). In Figure 3D-E, we show that our model is capable of predicting the overall structure of the cyclic peptide monomers with reasonable accuracy, including those with disulfide bonds.

**Figure 3.**
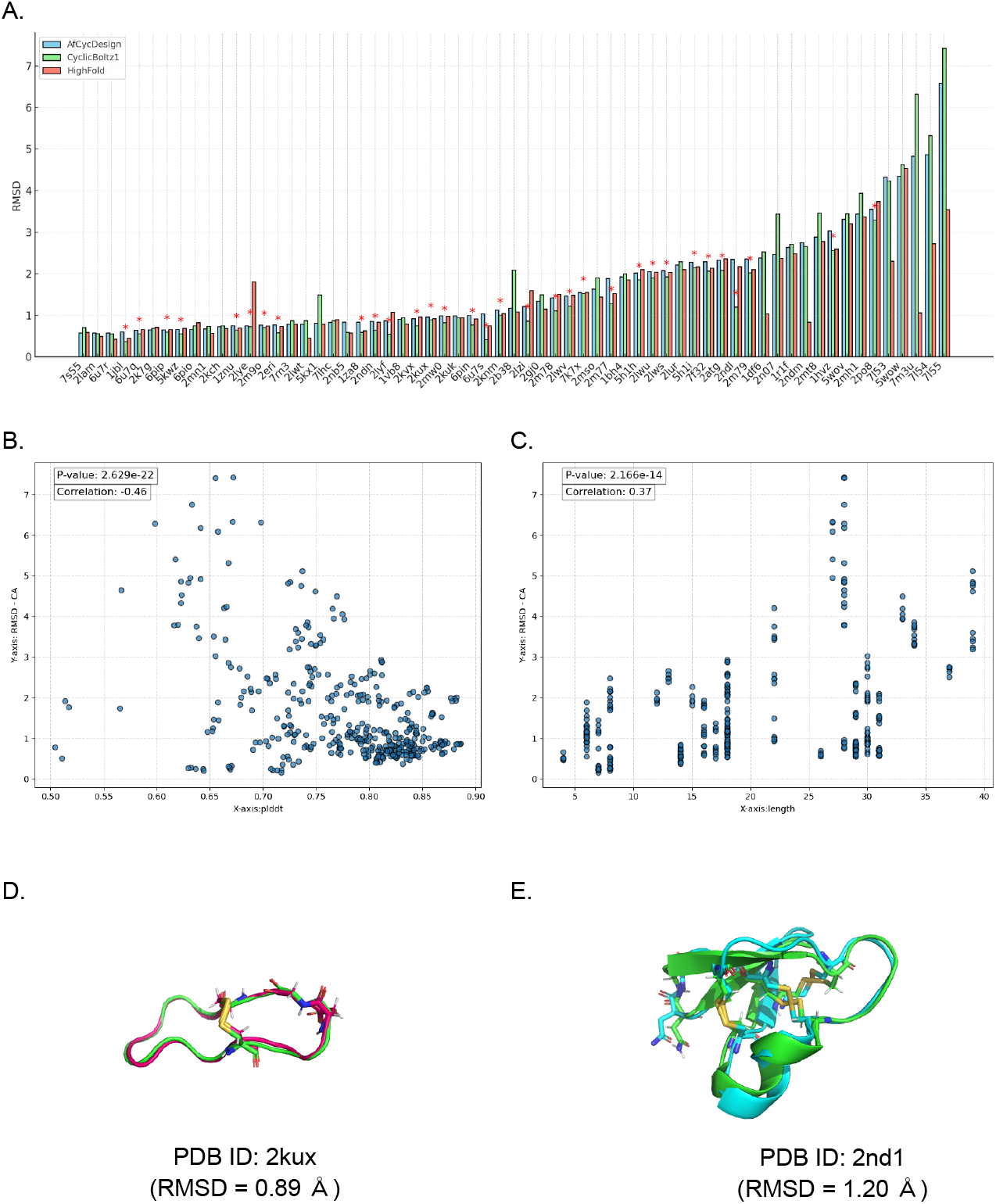
Cyclic peptide monomer structure prediction. **A**. C*α* RMSD distribution of CyclicBoltz1, HighFold, and AfCycDesign over the 63 benchmark test cases. The star indicates test cases where our model shows the lowest RMSD among all models. **B**. Scatter plot of C*α* RMSD and plDDT with Pearson correlation. **C**. Scatter plot of C*α* RMSD and peptide length with Pearson correlation. **D-E**.Visualization examples for prediction using CyclicBoltz1 with their corresponding C*α* RMSD.

### Prediction of Non-Canonical Amino Acid (ncAA)-containing Cyclic Peptide Monomers

Once we demonstrated that CyclicBoltz1-monomer was capable of generating cyclic peptides, we wanted to see if the model was able to predict structures of cyclic peptides with non-canonical amino acids (ncAAs). Using the HighFold2 dataset [16], we selected 17 structures with head-to-tail linkage to use as our benchmark against reported values from the HighFold2 paper. We generated 5 samples with CyclicBoltz1-monomer and took the top-1 sample for comparison against HighFold2 where they took the top-1 sample from the top-5 ranked predictions. In this case, we assessed both the C*α* RMSD as well as all-atom RMSD as the main metrics to capture the deviations in the sidechain prediction of the ncAAs during evaluations against the ground-truth structures.

We found that our model showed stronger overall performance compared to the reported HighFold2 performance in the prediction of monomers with modified residues. In Figure 4A-D, our model has higher accuracy than the HighFold2 model in 13 out of 17 test cases. Overall, when compared to ground truth structures, the predicted structures had a median C*α* RMSD of 1.798 Å with a median all-atom RMSD of 2.757 Å. Our results show significantly lower RMSDs when compared to the reported HighFold2 predictions, which had a median C*α* RMSD of 2.221 Å with a median all-atom RMSD of 3.307 Å. Similar to the previous monomer results, we found that there was a negative correlation between the RMSDs and the confidence metrics (confidence score and pLDDT) (Fig. 3C and Fig. 4E-F). From the various test cases, we observed that CyclicBoltz1-monomer can modify the standard amino acid given the corresponding Chemical Component Dictionary (CCD) code for the modified residue. We show that our model can modify the standard amino acid L-Proline to either D-Proline (PDB: 3wng) or 4-Hydroxy-Proline (PDB: 2j15). We show that adding the cyclic offset on top of the existing Boltz1 architecture allows structure prediction of cyclic peptide monomers containing ncAAs.

**Figure 4.**
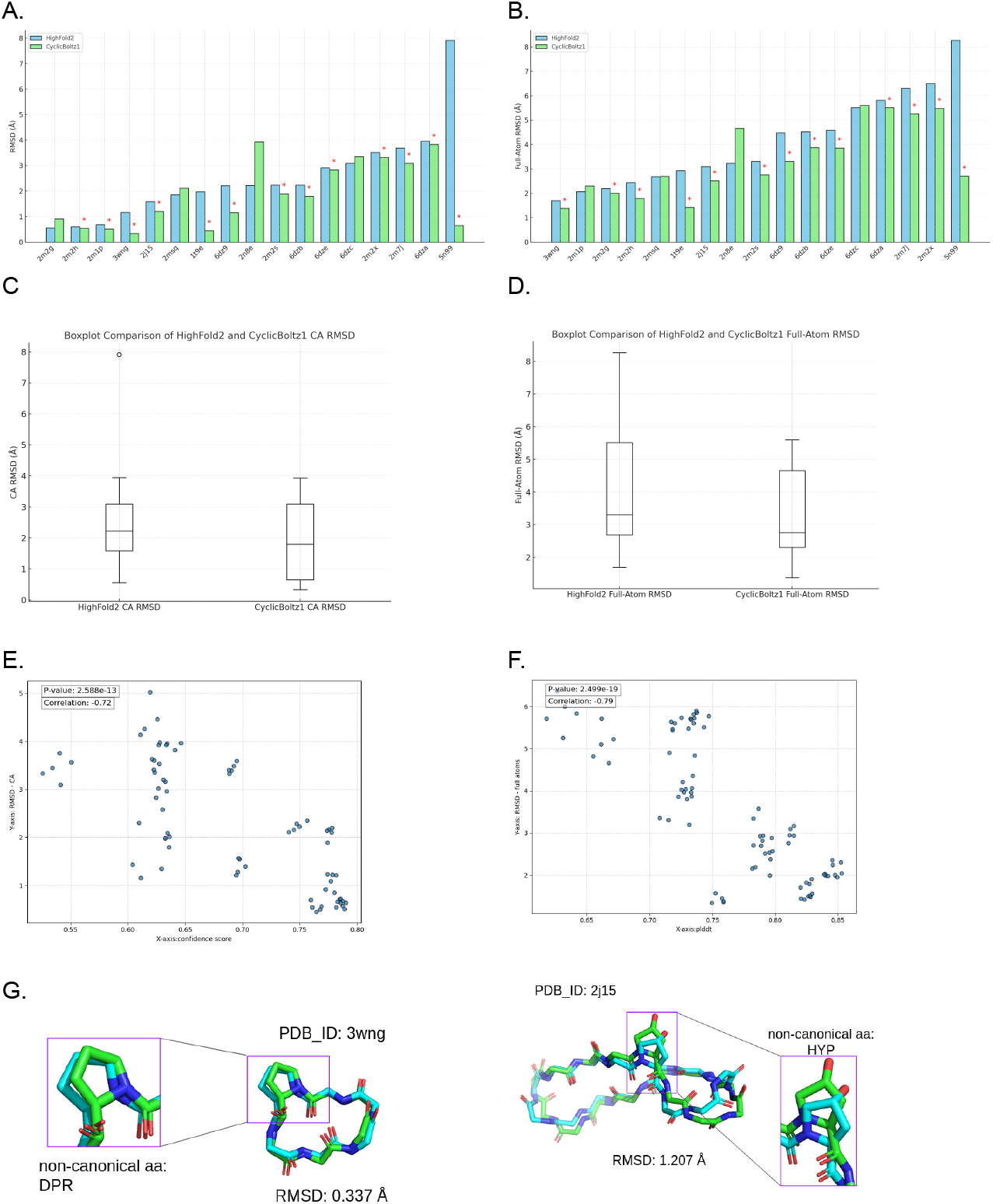
Cyclic peptide monomer structure prediction with ncAAs. **A**. Ca-RMSD distribution of CyclicBoltz1, and HighFold2 over the 17 non-canonical benchmark test cases. The star indicates test cases where our model shows the lowest RMSD among all models. **B**. Full-atom RMSD distribution of CyclicBoltz1, and HighFold2 over the 17 non-canonical benchmark test cases **C-D**. Boxplot comparison between CyclicBoltz1 and HighFold2 (C*α* and Full-atom RMSD). **E-F**. Scatter plot between C*α* RMSD and confidence metrics (pLDDT and confidence scores) with Pearson correlation **G**. Visualisation of predicted structures of cyclic peptide monomers containing ncAAs with their corresponding C*α* RMSD

### CyclicBoltz1-multimer Complex Prediction with Cyclic Peptide Ligands

Modeling only the cyclic peptide monomer, however, is insufficient for cyclic peptide binder design and evaluation. Often we wish to predict the structure of the entire complex containing both the target protein and corresponding cyclic peptide ligand to design potential binders. As such, we decided to test the complex structure prediction using CyclicBoltz1-multimer where we modified the cyclic encoding as shown in Fig 2D. To evaluate complex structure prediction, we utilized an external dataset [10] that was used to benchmark against HighFold_Multimer. We ran the HighFold_Multimer predictions as our control and evaluated both the ligand and receptor C*α* RMSD against the ground truth structures for both models.

We define the ligand RMSD as the C*α* RMSD of the ligand computed after alignment of the receptor with the ground truth receptor. Since the ligand C*α* RMSD is more relevant for the evaluation of complex prediction and docking of the ligand peptide, this was the primary metric considered for comparison between the two models. From the 15 test cases that we ran, we saw that our model showed stronger performance in 9 out of the 15 test cases (Fig. 5A) where we compared the top-1 prediction for each model. In this case (Fig. 5B-C), we see a moderate correlation between the C*α* RMSD and the confidence metrics (pLDDT and confidence score) likely due to the combined diversity of multiple peptide and protein sequences and structures. In the majority of test cases, we found that our model was able to place the generated cyclic peptide in the same pocket as the ground-truth structures (Fig. 5D-F).

**Figure 5.**
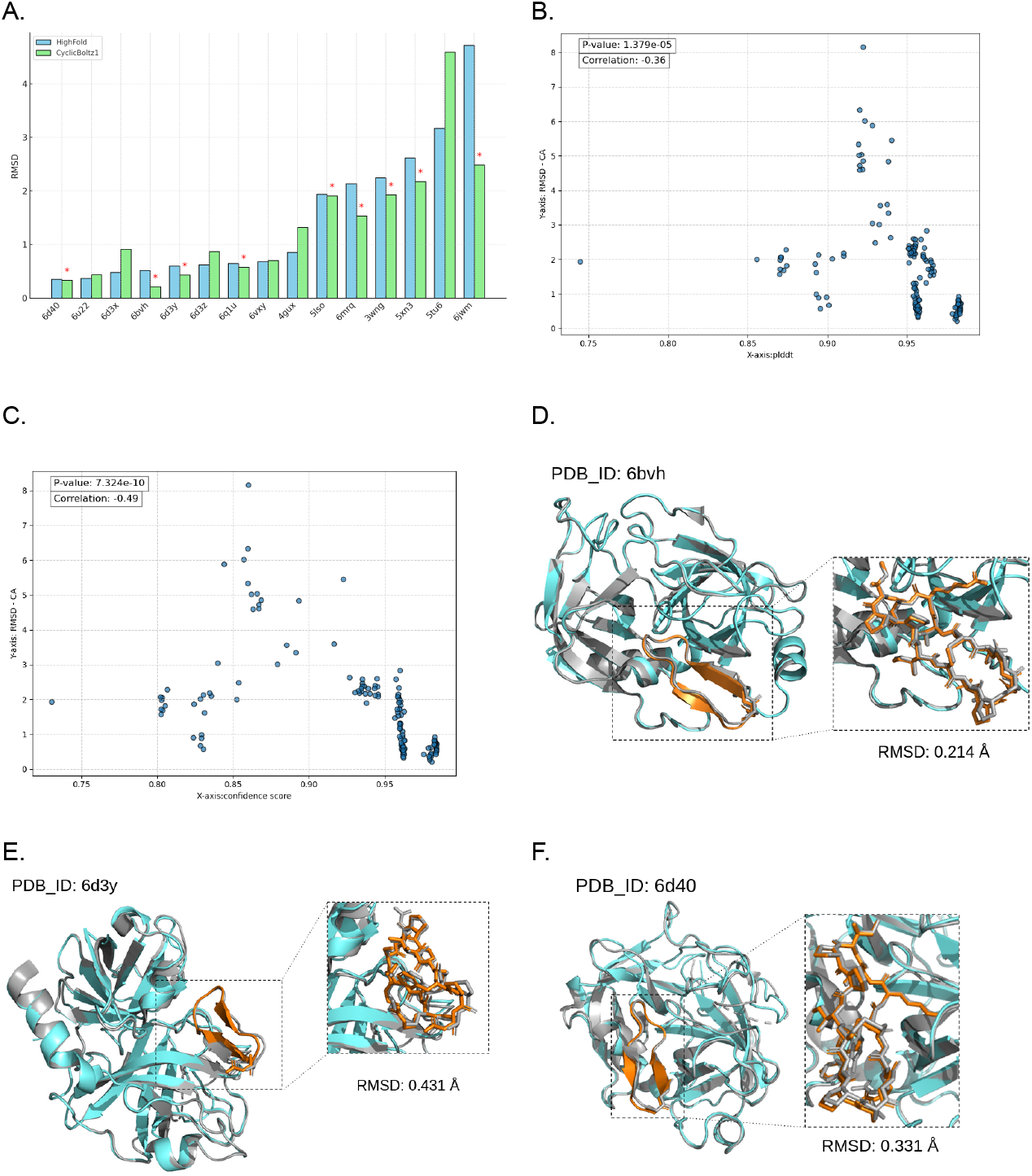
Multimer Benchmark Results. **A**. Ligand C*α* RMSD distribution of CyclicBoltz1, and HighFold over the 15 benchmark test cases. The star indicates those test cases where our model shows the lowest RMSD among all models. **B-C**. Scatter plot between C*α* RMSD and confidence metrics (pLDDT and confidence score) with Pearson correlation **D-F**. Visualization examples for prediction using CyclicBoltz1 with their corresponding C*α* RMSD.

## Discussions

By incorporating the cyclic offset encoding, achieved through modifications to the model’s relative positional encoding matrix, CyclicBoltz1 enables precise predictions of cyclic peptides while facilitating multi-scale modeling of peptide molecules. Testing demonstrated outstanding performance on both independent cyclic peptide test sets and modified peptide datasets. CyclicBoltz1 can predict cyclic peptide monomer structures with ncAAs as well as multimeric complexes with cyclic peptide ligands. Overall, we were able to show comparable results to existing cyclic peptide structure prediction models.

We are currently in the process of testing our model against a diverse range of datasets containing different modifications and sequences. Thus far, we’ve mainly tested against the HighFold and HighFold2 datasets and aim to benchmark against more complex and monomeric structures. Concurrently, we are expanding our structure prediction model to include binder design as shown in Fig. 1C. Further experimental testing will need to be done in the future to validate our results.

In conclusion, our method bypasses the scarcity of cyclic peptide structural data by leveraging existing pre-trained models developed for protein and peptide structure prediction and design that enable residue modifications. We’ve shown that applying a cyclic offset to AlphaFold3 enables structure prediction for cyclic peptides which is especially useful for modeling cyclic peptides containing ncAAs. As such, we have developed a robust tool for advancing cyclic peptide structure prediction and design.

## Supporting information

Supplementary Algorithm 1

## References

[1] Wang, L. (2022). Therapeutic peptides: Current applications and future directions. Signal Transduction and Targeted Therapy.

[2] Fosgerau, K., & Hoffmann, T. (2015). Peptide therapeutics: current status and future directions. Drug discovery today, 20(1), 122–128.

[3] Driggers, E. M., Hale, S. P., Lee, J., & Terrett, N. K. (2008). The exploration of macrocycles for drug discovery--an underexploited structural class. Nature reviews. Drug discovery, 7(7), 608–624. 10.1038/nrd2590

[4] Evans, R., O’Neill, M., Pritzel, A., Antropova, N., Senior, A., Green, T., Žídek, A., Bates, R., Blackwell, S., Yim, J., Ronneberger, O., Bodenstein, S., Zielinski, M., Bridgland, A., Potapenko, A., Cowie, A., Tunyasuvunakool, K., Jain, R., Clancy, E., … Hassabis, D. (2021). Protein complex prediction with AlphaFold-Multimer. bioRxiv, 2021.10.04.463034. 10.1101/2021.10.04.463034

[5] Abramson, J., Adler, J., Dunger, J., Evans, R., Green, T., Pritzel, A., … & Jumper, J. M. (2024). Accurate structure prediction of biomolecular interactions with AlphaFold 3. Nature, 1–3.

[6] Krishna, R., Wang, J., Ahern, W., Sturmfels, P., Venkatesh, P., Kalvet, I., Lee, G. R., Morey-Burrows, F. S., Anishchenko, I., Humphreys, I. R., McHugh, R., Vafeados, D., Li, X., Sutherland, G. A., Hitchcock, A., Hunter, C. N., Kang, A., Brackenbrough, E., Bera, A. K., Baek, M., … Baker, D. (2024). Generalized biomolecular modeling and design with RoseTTAFold All-Atom. Science (New York, N.Y.), 384(6693), eadl2528. 10.1126/science.adl2528

[7] Jumper, J., Evans, R., Pritzel, A., Green, T., Figurnov, M., Ronneberger, O., … & Hassabis, D. (2021). Highly accurate protein structure prediction with AlphaFold. nature, 596(7873), 583–589.

[8] Baek, M., DiMaio, F., Anishchenko, I., Dauparas, J., Ovchinnikov, S., Lee, G. R., … & Baker, D. (2021). Accurate prediction of protein structures and interactions using a three-track neural network. Science, 373(6557), 871–876.

[9] Rettie, S. A., Campbell, K. V., Bera, A. K., Kang, A., Kozlov, S., De La Cruz, J., Adebomi, V., Zhou, G., DiMaio, F., Ovchinnikov, S., & Bhardwaj, G. (2023). Cyclic peptide structure prediction and design using AlphaFold. bioRxiv, 2023.02.25.529956. 10.1101/2023.02.25.529956

[10] Zhang, C., Zhang, C., Shang, T., Zhu, N., Wu, X., & Duan, H. (2024). HighFold: accurately predicting structures of cyclic peptides and complexes with head-to-tail and disulfide bridge constraints. Briefings in bioinformatics, 25(3), bbae215. 10.1093/bib/bbae215

[11] Rettie, S. A., Juergens, D., Adebomi, V., Bueso, Y. F., Zhao, Q., Leveille, A. N., Liu, A., Bera, A. K., Wilms, J. A., Üffing, A., Kang, A., Brackenbrough, E., Lamb, M., Gerben, S. R., Murray, A., Levine, P. M., Schneider, M., Vasireddy, V., Ovchinnikov, S., … Bhardwaj, G. (2024). Accurate de novo design of high-affinity protein binding macrocycles using deep learning. bioRxiv, 2024.11.18.622547. 10.1101/2024.11.18.622547

[12] Watson, J. L., Juergens, D., Bennett, N. R., Trippe, B. L., Yim, J., Eisenach, H. E., Ahern, W., Borst, A. J., Ragotte, R. J., Milles, L. F., Wicky, B. I. M., Hanikel, N., Pellock, S. J., Courbet, A., Sheffler, W., Wang, J., Venkatesh, P., Sappington, I., Torres, S. V., Lauko, A., … Baker, D. (2023). De novo design of protein structure and function with RFdiffusion. Nature, 620(7976), 1089–1100. 10.1038/s41586-023-06415-8

[13] Berman, H. M., Westbrook, J., Feng, Z., Gilliland, G., Bhat, T. N., Weissig, H., Shindyalov, I. N., & Bourne, P. E. (2000). The Protein Data Bank. Nucleic acids research, 28(1), 235–242. 10.1093/nar/28.1.235

[14] Wohlwend, J., Corso, G., Passaro, S., Reveiz, M., Leidal, K., Swiderski, W., Portnoi, T., Chinn, I., Silterra, J., Jaakkola, T., & Barzilay, R. (2024). Boltz-1 Democratizing Biomolecular Interaction Modeling. bioRxiv, 2024.11.19.624167. 10.1101/2024.11.19.624167

[15] Costa, L., Sousa, E., & Fernandes, C. (2023). Cyclic Peptides in Pipeline: What Future for These Great Molecules? Pharmaceuticals, 16(7), 996. 10.3390/ph16070996

[16] Zhu, C., Cao, S., Shang, T., Guo, J., Su, A., Li, C., & Duan, H. (2025). Predicting the structures of cyclic peptides containing unnatural amino acids by HighFold2. bioRxiv, 2025.01.16.633493. 10.1101/2025.01.16.633493

[17] Du, Y., Li, L., Zheng, Y., Liu, J., Gong, J., Qiu, Z., … & Huo, Y. X. (2022). Incorporation of non-canonical amino acids into antimicrobial peptides: advances, challenges, and perspectives. Applied and Environmental Microbiology, 88(23), e01617–22.

[18] Miao, J., Descoteaux, M. L., & Lin, Y. S. (2021). Structure prediction of cyclic peptides by molecular dynamics+ machine learning. Chemical science, 12(44), 14927–14936.

[19] Charitou, V., Van Keulen, S. C., & Bonvin, A. M. (2022). Cyclization and docking protocol for cyclic peptide–protein modeling using HADDOCK2. 4. Journal of chemical theory and computation, 18(6), 4027–4040.

[20] Karami, Y., Murail, S., Giribaldi, J., Lefranc, B., Defontaine, F., Lesouhaitier, O., … & Tufféry, P. (2022). Head-to-tail peptide cyclization: new directions and application to urotensin II and Nrf2. bioRxiv, 2022-01.

