## Supplementary Algorithm 1 for "CyclicBoltz1, fast and accurately predicting structures of cyclic peptides and complexes containing non-canonical amino acids using AlphaFold 3 Framework"

### Modified cyclic peptide matrix for ncAAs

Relative positional encoding features capture index distance information between amino acids in linear structures. AfCycDesign, HighFold, and RFpeptides adjusted their positional encodings to ensure that N- and C-terminal residues remain at a fixed index distance of 1 using a cyclic offset matrix. However, previous cyclic offset encoding strategies operated only at the residue level and failed in Boltz1 due to structural modifications—specifically, AlphaFold3's introduction of atom token representations into relative encoding features for modified residues (ncAAs). To address this, we modified the cyclic encoding matrix using unique indexing to ensure consistent relative encodings per residue index (Algorithm 1). Given Boltz1's atom-token-based encoding, the resulting relative positional encoding matrix reflects this structure, as shown in Figure 2C, where blue rows indicate standard residue tokens and yellow rows correspond to modified residue atom tokens.

**Algorithm 1 Cyclic Offset with non-canonical Amino Acids**

```
function cyclic_offset with_modified_aa(residue_index)
    unique_indices, inverse_indices = unique(residue_index)
    #Get unique indices and inverse indices
    L = size(unique_indices)
    # Logical residue length
    i = arange(L)
    ij = stack([i, i + L], −1)
    of f_set = i[:, None] − i[None, :]  # Pairwise index differences
    c_off_set = abs(ij[:, None, :, None] − ij[None, :, None, :])
    c_off_set = min(c_offset, −1)  # Minimum cyclic distance
    a = c off set < abs(offset)
    c_offset[a] = −c_offset[a]  # Negate when needed
    expanded_c_of f_set = c_of fset[inverse indices][:, inverse indices]
    # Restore original shape
    return expanded_c_offset × sign(offset[inverse indices][:, inverse indices])
end function
```
